# MoB (Measurement of Biodiversity): a method to separate the scale-dependent effects of species abundance distribution, density, and aggregation on diversity change

**DOI:** 10.1101/244103

**Authors:** Daniel J. McGlinn, Xiao Xiao, Felix May, Nicholas J. Gotelli, Thore Engel, Shane A. Blowes, Tiffany M. Knight, Oliver Purschke, Jonathan M. Chase, Brian J. McGill

## Abstract

1. Little consensus has emerged regarding how proximate and ultimate drivers such as productivity, disturbance, and temperature may affect species richness and other aspects of biodiversity. Part of the confusion is that most studies examine species richness at a single spatial scale and ignore how the underlying components of species richness can vary with spatial scale.
2. We provide an approach for the measurement of biodiversity (MoB) that decomposes changes in species rarefaction curves into proximate components attributed to: 1) the species abundance distribution, 2) density of individuals, and 3) the spatial arrangement of individuals. We decompose species richness by comparing spatial and nonspatial sample- and individual-based species rarefaction curves that differentially capture the influence of these components to estimate the relative importance of each in driving patterns of species richness change.
3. We tested the validity of our method on simulated data, and we demonstrate it on empirical data on plant species richness in invaded and uninvaded woodlands. We integrated these methods into a new R package (`mobr`).
4. The metrics that `mobr` provides will allow ecologists to move beyond comparisons of species richness in response to ecological drivers at a single spatial scale towards a dissection of the proximate components that determine species richness across scales.

## Introduction

Species richness – the number of species co-occurring in a specified area – is one of the most widely-used biodiversity metrics. However, ecologists often struggle to understand the mechanistic drivers of richness, in part because multiple ecological processes can yield qualitatively similar effects on species richness (Chase and Leibold 2002, Leibold and Chase 2017). For example, high species richness in a local community can be maintained either by species partitioning niche space to reduce interspecific competition (Tilman 1994), or by a balance between immigration and stochastic local extinction (Hubbell 2001). Similarly, high species richness in the tropics has been attributed to numerous mechanisms such as higher productivity supporting more individuals, higher speciation rates, and longer evolutionary time since disturbance (Rosenzweig 1995).

Although species richness is a single metric that can be measured at a particular grain size or spatial scale, it summarizes the underlying biodiversity information that is contained in the individual organisms, each of which are assigned to a particular species, Operational Taxonomic Unit, or other taxonomic grouping. Variation in species richness can be decomposed into three components (He and Legendre 2002, McGill 2010, 2011a): 1) the number and relative proportion of species in the regional source pool (i.e., the species abundance distribution, SAD), 2) the number of individuals per plot (i.e., density), and 3) the spatial distribution of individuals that belong to the same species (i.e., spatial aggregation). Changes in species richness from one place to another (or one time to another) may reflect one or a combination of all three components changing simultaneously (Chase and Knight 2013). Although sampling intensity and detection error influence these observations, they are also strongly influenced by experimental or observational treatments that ultimately drive the patterns of observed species richness. Thus, it is critical that we look beyond richness as a single metric, and develop methods to disentangle its underlying components that have more mechanistic links to processes (e.g., Vellend 2016).

While there are other mathematically valid decompositions of species richness and its change, the three components above (density, SAD and aggregation) are well-studied properties of ecological systems, and provide insights into mechanisms behind changes in richness and community structure (Harte et al. 2008, Supp et al. 2012, McGlinn et al. 2015). We note that the SAD and N components are also sometimes referred to as the column and row sums respectively of the abundance-based community matrix (e.g., Ulrich and Gotelli 2010).

Local richness is influenced by the shape of the regional SAD which reflects the degree to which common species dominate the individuals observed in a region, and on the total number of species in the pool. Local communities that are part of a more even regional SAD (i.e., most species having similar abundances) will have high values of local richness because it is more likely that the individuals sampled will represent different species. Local communities that are part of regions with a more uneven SAD (e.g., most individuals are a single species) will have low values of local richness because it is more likely that the individuals sampled will be the same common species (He and Legendre 2002, McGlinn and Palmer 2009). The richness of the regional species pool can also influence local richness. As regional species richness increases, local richness will also increase if the local community is even a partly random subsample of the species in the regional pool. Because the regional species pool is never fully observed, the two sub-components – the shape of the SAD and the size of the regional species pool – cannot be completed disentangled. Thus, we group them together, as the SAD effect on local richness.

The number of individuals in the local community directly affects richness due to the sampling effect (the “More Individuals Hypothesis”, Srivastava and Lawton 1998). As more individuals are sampled from the regional pool, species richness is bound to increase via sampling. This effect will be strongest at fine spatial scales; however, even at larger spatial scales, it never truly goes to zero (Palmer and van der Maarel 1995, Palmer et al. 2008).

The spatial arrangement of individuals in a landscape is rarely random. Instead most individuals are spatially clustered or aggregated in some way, with neighboring individuals more likely belonging to the same species. As of clusters of only a few individuals within species become more spatially clustered, local diversity will decrease because the local community or sample is likely to consist of clusters of only a few species (He and Legendre 2002, Karlson et al. 2007, Chiarucci et al. 2009, Collins and Simberloff 2009).

Traditionally, individual-based rarefaction has been used to control for the effect of numbers of individuals on richness comparisons (Hurlbert 1971, Simberloff 1972, Gotelli and Colwell 2001), but few methods exist (e.g., Cayuela et al. 2015) for decomposing the effects of SADs and spatial aggregation on species richness. Because species richness depends intimately on the spatial and temporal scale of sampling, the relative contributions of the three components discussed above are also likely to change with scale. Spatial scale can be represented both by the numbers of samples (plots) taken, and by number of individuals within those plots, which scale linearly with area (McGill 2011a, see Scale and sampling effort in Methods, also see Supplement S7). Below, we will demonstrate that this generalized view of spatial scale allows us to synthesize the information provided by three different types of rarefaction curves: (1) spatially constrained, sample-based rarefaction; (2) non-spatial, sample-based rarefaction; and (3) (non spatial) individual-based rarefaction. Constructing these different curves allows us to parse the relative contributions of the three components of richness and how those contributions potentially change with spatial scale. Specifically, we develop a framework that provides a series of sequential analyses for estimating and testing the effects of the SAD, individual density, and spatial aggregation on changes in species richness across scales. We have implemented these methods in a freely available R package `mobr` (https://github.com/MoBiodiv/mobr)

## Materials and Methods

### Method Overview

Our method targets data collected in standardized sampling units such as quadrats, plots, transects, net sweeps, or pit falls of constant area or sampling effort (we refer to these as “plots”) that are assigned to treatments. We use the term treatment here generically to refer to manipulative treatments or to groups within an observational study (e.g., invaded vs uninvaded plots). It is critical that the treatments have identical grain (i.e., area of the plots) and similar plot spatial arrangements across similar extents. If the sampling design differs between the treatments then treatment effects will be obscured by scale-mismatches; nevertheless, this can often be remedied post-hoc through various types of standardization.

In an experimental study, each plot is assigned to a treatment. In an observational study, each plot is assigned to a categorical grouping variable(s). For this typical experimental/sampling design, our method provides two key outputs: 1) the observed change in richness between treatments and the relative contribution of the different components affecting richness (SAD, density, and spatial aggregation) to those changes and 2) how species richness and its decomposition change with spatial scale. We propose two complementary ways to view scale-dependent shifts in species richness and its components: a simple-to-interpret two-scale analysis and a more informative but necessarily more complex multi-scale analysis.

The two-scale analysis provides a big-picture view of the changes between the treatments by focusing exclusively on the *α* (plot-level) and *γ* (across all plots) spatial scales. It provides diagnostics for whether species richness and its components differ between treatments at the two scales. The multi-scale scale analysis expands the two-scale analysis by taking advantage of three distinct types of rarefaction curves: 1) spatially constrained, sample-based rarefaction (Chiarucci et al. 2009), where the order in which plots are sampled depends on their spatial proximity (these are referred to as species accumulation curves in Gotelli and Colwell 2001); 2) the non-spatial, sample-based rarefaction, where individuals are randomly shuffled across plots within a treatment while maintaining average density in the plots; and 3) the individual-based rarefaction curve where again individuals are randomly shuffled across plots within a treatment, but in this case average plot density is not maintained (Hurlbert 1971, Gotelli and Colwell 2001). The differences between these curves are used to isolate the effects of the SAD, density of individuals, and spatial aggregation on richness and document how these effects change as a function of scale.

### Scale and sampling effort

Grain or the scale of interest (focus *sensu* Scheiner et al. 2000) can be varied by considering different numbers of individuals or plots (i.e., the sampling effort) from the α-scale (a single plot) to the *γ*-scale (all plots in a treatment). It is possible to interchange numbers of individuals. numbers of plots, and spatial area because the average number of individuals accumulated scales linearly with the number of plots or area (McGill 2011a, Supplement S7). This is true when considering both spatially contiguous and non-contiguous sampling designs.

Theoretical treatments of the species-area relationship established the connection between the number of individuals and sampled area (Arrhenius 1921, Williams 1943, Hubbell 2001, Harte 2011). The MoB framework uses this connection to synthesize the information provided by the three types of rarefaction curves which differ only in how they define sampling effort (i.e., scale).

### Mathematical nomenclature

Consider *T* = 2 treatments, with *K* replicated plots per treatment (See supplemental Table S1.1). Within each plot, we have measured the abundance of each species, which can be denoted by *n_t,k,s_*, where *t* = 1, 2 for treatment, *k* = 1, 2, … *K* for plot number within the treatment, and *s* = 1, 2, … *S* for species identity, with a total of *S* species recorded among all plots and treatments. The experimental design does not necessarily have to be balanced (i.e., *K* can differ between treatments) so long as the spatial extent is similar between the treatments. However, it is important to note that inferences can only be made over a common number of individuals or samples, so large differences in sampling effort will decrease the range of scales useful for assessing treatment effects (but this can be remedied by standardization of sampling among treatments post-hoc). For simplicity of notation we describe the case of a balanced design here. *S_t,k_* is the number of species observed in plot *k* in treatment *t* (i.e., number of species with *n_t,k,s_* > 0), and *N_t,k_* is the number of individuals observed in plot *k* in treatment *t* (i.e., *N_t,k_* = Σ_*s*_*n_t,k,s_*). The spatial coordinates of each plot *k* in treatment *t* are *x_t,k_* and *y_t,k_*. We focus on spatial patterns but our framework also applies analogously to samples distributed through time.

For clarity, we focus here on a single-factor design with two (or more) categorical treatment levels. The method can and will be extended to accommodate crossed designs and regression-style continuous treatments, which we describe in the Discussion and Supplement S6.

### Two-scale analysis

The two-scale analysis provides a simple decomposition of species richness while still emphasizing the three components and change with spatial scale. In the two-scale analysis, we compare observed species richness in each treatment and several other summary statistics at the *α* and *γ* scales (Supplemental Table S2). The summary statistics were chosen to represent the most informative aspects of individual-based rarefaction curves (Fig. 1). Individual-based rarefaction curves plot the expected species richness *S_n_* against the number of individuals when individuals are randomly drawn from the sample at the *α* or *γ* scales. The curve can be calculated precisely using the hypergeometric sampling formula given the SAD (*n_t,k,s_* at the *α*-scale, *n_t,+,s_* at the *γ*-scale) (Hurlbert 1971).

**Figure 1.**
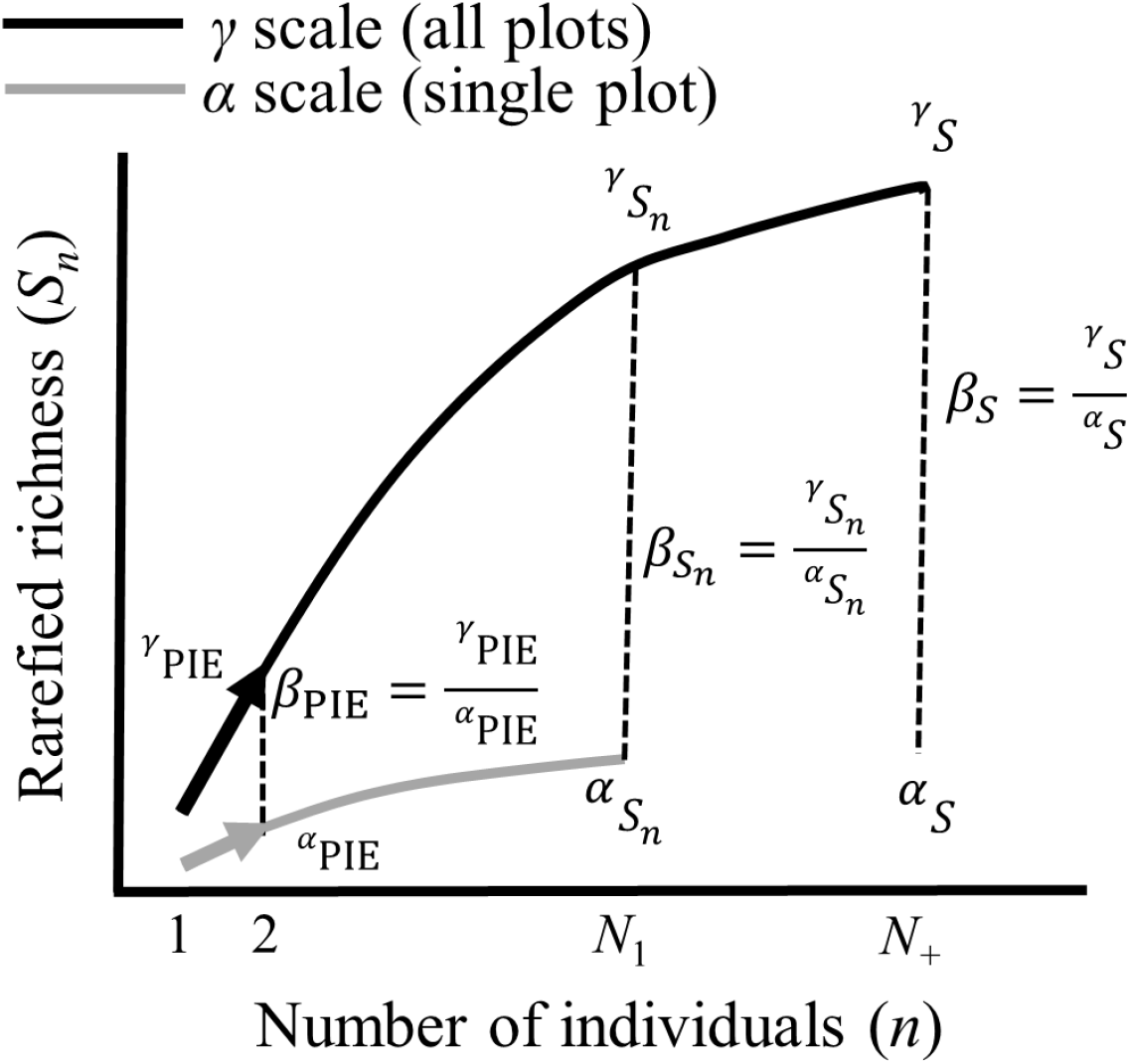
Illustration of how the key biodiversity metrics are derived from the individual-based rarefaction curves constructed at the *α* (i.e., single plot, grey line) and *γ* (i.e., all plots, black line) scales. The curves are rarefied richness derived from the randomly sampling individuals from either the SAD of one or all plots. See Supplemental Table S1.2 for definitions of metrics including ones not illustrated.

We show how several widely-used diversity metrics are represented along the individual rarefaction curve, corresponding to *α* and *γ* scales (Fig. 1, see Supplement S1 for detailed metric description, Chase et al. 2018 provided a more detailed description and justification of the two-scale analysis). The total number of individuals within a plot (*N_t,k_*) or within a treatment (*N_t,+_*) determines the endpoint of the rarefaction curves. Rarefied richness (Sn) controls richness comparisons for differences in individual density between treatments because it is the expected number of species for a random draw of *n* individuals ranging from 1 to *N*. To compute *S_n_* at the *α*-scale, we set *n* to the minimum total number of individuals across all samples in both treatments with a hard minimum of 5, and at the *γ* scale we multiplied this *n* value by the number of samples within a treatment (i.e., *K*). The SAD component is complex, but aspects of it can be understood by evaluating the relative abundances of species (e.g., via an evenness metric). Here, we use the probability of interspecific encounter (PIE) (Hurlbert 1971) as a measure of evenness that captures differences in the relative abundances of the most common species; rare species can be examined via a number of different metrics many of which are highly correlated with *S* (McGill 2011b). Importantly, PIE represents the slope at the base of the individual-based rarefaction curve (Olszweski 2004) (Fig. 1). For analyses, we convert PIE into an asymptotic effective numbers of species (S_PIE_) so that it can be more easily interpreted as a metric of diversity (Supplement S2). Whittaker’ s multiplicative *β*-diversity metrics for *S, S*_PIE_, and *S_n_* reflect the degree of turnover between the *α*- and *γ*-scales. *β_Sn_* is computed at the same *n* value at the *α*- and *γ*-scales (Fig. 1) to control for differences in individual density and SADs. Therefore, *β_Sn_* provides a means of isolating the effect of intraspecific aggregation on turnover patterns while controlling for SAD and *N* effects (Chase et al. 2018).

Comparison of these summary statistics between treatments identifies whether the treatments have a significant effect on richness at these two scales, and if they do, the potential proximate component(s) of the change (Chase et al. 2018). A treatment difference in *N* implies that differences in richness may be a result of treatments changing the density of individuals. Differences in *S*_PIE_ imply that change in the shape of the SAD may contribute to the change in richness, with *S*_PIE_ being most sensitive to changes in abundant species and *S* being most sensitive to changes in number of rare species. Differences in *β*-diversity metrics may be due to differences in any of the three components: SAD, *N*, or aggregation. *β_Sn_* is unique in that it attempts to control for SAD and *N* effects on species turnover to isolate the signal of intraspecific aggregation (Chase et al. 2018).

The treatment effect on these metrics can be visually examined with boxplots (see Empirical example section) at the *α*-scale and with single points at the pooled *γ*-scale (unless there is replication at the *γ*-scale as well). Quantitative comparison of the metrics can be made with t-tests (ANOVAs for more than two treatments) or, for highly skewed data, nonparametric tests such as Mann-Whitney U test (Kruskal-Wallis for more than two treatments).

We provide a non-parametric, randomization test where the null expectation of each metric is established by randomly shuffling the plots between the treatments, and recalculating the metrics for each reshuffle. The significance of the differences between treatments can then be evaluated by comparing the observed test statistic to the null expectation when the treatment IDs are randomly shuffled across the plots (Legendre and Legendre 2012). When more than two groups are compared, the test examines the overall group effect rather than specific group differences. At the *α*-scale where there are replicate plots to summarize over, we use the ANOVA *F*-statistic as our test statistic (Legendre and Legendre 2012), and at the *γ*-scale in which we only have a single value for each treatment (and therefore cannot use the *F*-statistic) the test statistic is the absolute difference between the treatments (if more than two treatments are considered then it is the average of the absolute differences, 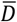). We use 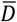 as a measure of effect size at both scales.

Note that *N_t,k_* and *N_t,+_* give the same information, because one scales linearly with the other by a constant (i.e., *N_t,+_* is equal to *N_t,k_* multiplied by the number of plots *K* within treatment). However, the other metrics (*S, S_n_*, and *S*_PIE_) are not directly additive across scales. Evaluation of these metrics at different scales may yield different insights for the treatments, sometimes even in opposite directions (Chase et al. 2018). However, complex scale-dependence may require comparison of entire rarefaction curves (rather than their two-scale summary statistics) to understand how differences in community structure change continuously across a range of spatial scales.

### Multiscale analysis

While the two-scale analysis provides a useful tool with familiar methods, it ignores that scale itself is not discrete, but rather is a continuum. Such a discrete scale perspective can thus only provide a minimal view of the scaling relationships of treatment differences. In this section, we develop a method to examine the components of change across a continuum of spatial scale. We define spatial scale by the amount of sampling effort (i.e., the number of individuals and/or the number of plots sampled; see Scale and sampling effort subsection)

### The three curves

The key innovation is to use three distinct types of rarefaction curves that capture different components of community structure. By a carefully sequenced analysis, it is possible to tease apart the effects of SAD shape, of changes in density of individuals (*N*), and of spatial aggregation across a continuum of spatial scale. The three types of curves are summarized in Table 1. Fig. 2 shows how they are constructed graphically.

**Figure 2.**
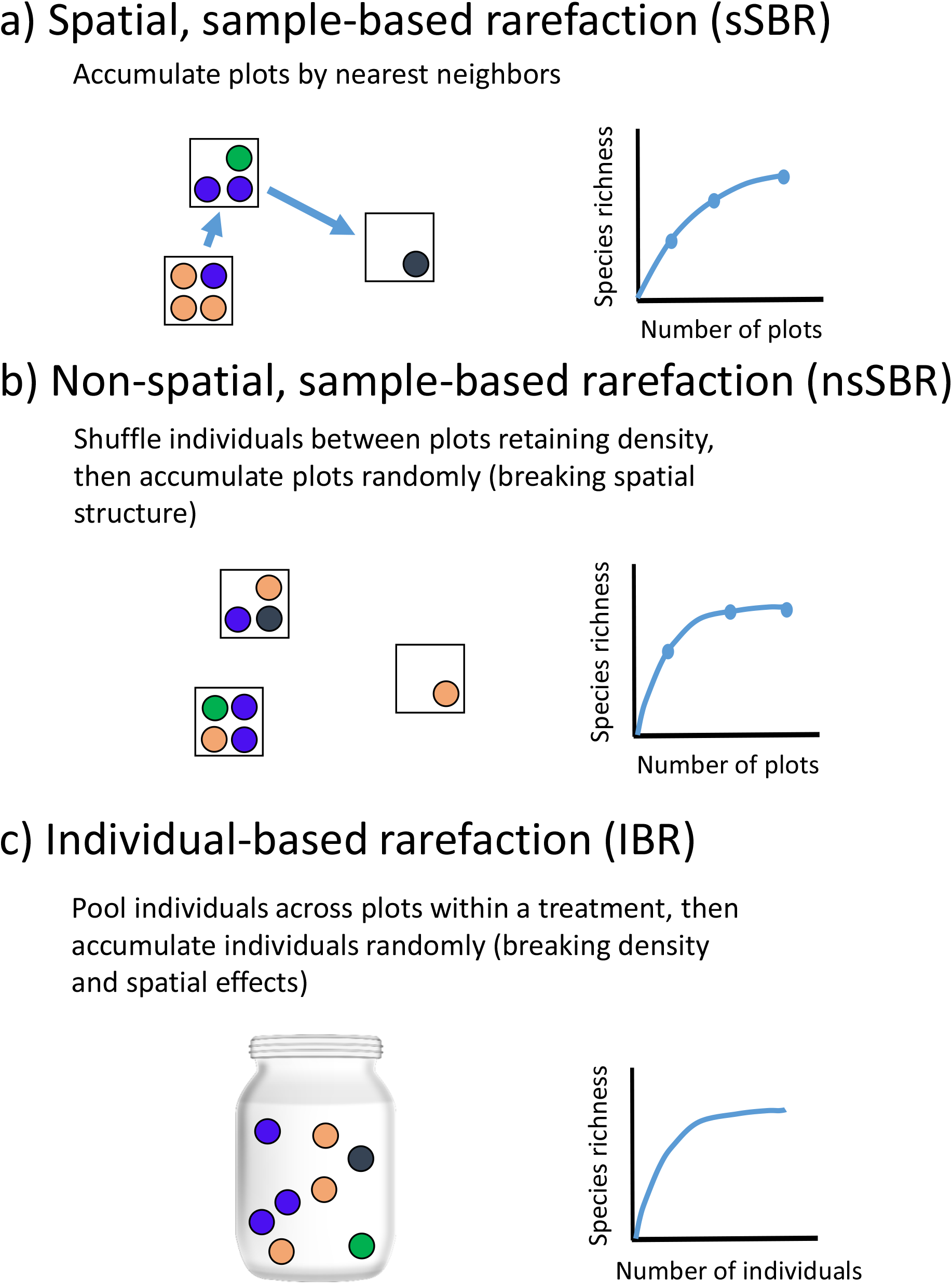
Illustration of how the three rarefaction curves are constructed. Circles of different colors represent individuals of different species. See Table 1 for detailed description of each rarefaction curve.

**Table 1.** Summary of three types of species rarefaction curves. For treatment *t*, 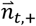 is the vector of species abundances, 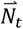 is the vector of plot abundances, and 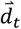 is the vector of distances between plots.

| Curve Name | Acronym | Notation | Method for accumulation | Interpretation |
| --- | --- | --- | --- | --- |
| Spatial,<br>sample-based<br>rarefaction<br>curve | sSBR | $E[S_t \vert k, \vec{n}_{t,+}, \vec{N}_t, \vec{d}_t]$ | Spatially explicit sampling in which the most proximate plots to a focal plot are accumulated first. All possible focal plots are considered and the resulting curves are averaged over. | The sSBR includes all information in the data including effect of SAD, effect of density of individuals, and effect of spatial aggregation. |
| Nonspatial,<br>sample-based<br>rarefaction<br>curve | nsSBR | $E[S_t \vert k, \vec{n}_{t,+}, \vec{N}_t]$ | Random sampling of $k$ plots after removing intraspecific spatial aggregation by randomly shuffling individuals across plots while maintaining average plot-level abundance ($\overline{N_{t,k}}$) and the treatment-level SAD ($n_{t,+s} = \sum_k n_{t,k,s}$). We use an analytical extension of the hypergeometric distribution demonstrating this curve is a rescaling of the IBR based on the ratio: (average density across treatments) / (average density of treatment of interest) | The nsSBR reflects both the shape of the SAD and the difference in density between the treatments. If density between the two treatments is identical then this curve converges on the IBR. |
| Individual-based<br>rarefaction<br>curve (IBR) | IBR | $E[S_t \vert N, \vec{n}_{t,+}]$ | Random sampling of $N$ individuals from the observed SAD ($\vec{n}_{t,+}$) without replacement. | By randomly shuffling individuals with no reference to plot density, all spatial and density effects are removed. Only the effect of the SAD remains. |

The first curve is the spatial, sample-based rarefaction (sSBR) (spatially-constrained rarefaction Chiarucci *et al*. 2009). It is constructed by accumulating plots sampled within a treatment based on their spatial position such that the most proximate plots are collected first.

One can think of this as starting with a target plot and then expanding a circle centered on the target plot until one additional plot is added, then expanding the circle until another plot is added, etc. In practice, every plot is used as the starting target plot and the resulting curves are averaged to give a smoother curve. If two or more plots are of equal distance to the target plot, they are accumulated in random order.

The second curve is the non-spatial, sample-based rarefaction curve (nsSBR, Supplement S3). It is constructed by randomly sampling plots within a treatment, but in which the individuals in the plots have first been randomly shuffled among the plots within a treatment, while maintaining the plot-level average abundance 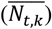 and the treatment-level SAD 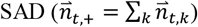. Note that this rarefaction curve is distinct from the traditional “sample-based rarefaction curve” (Gotelli and Colwell 2001), in which plots are randomly shuffled to build the curve but individuals within a plot are preserved (and consequently any within-plot spatial aggregation is retained). The nsSBR contains much of the same information as the sSBR (plot density and SAD), but it has nullified any signal due to species spatial aggregation.

The third curve is the familiar individual-based species rarefaction (IBR) (Hurlbert 1971, Gotelli and Colwell 2001). It is constructed by first pooling individuals across all plots within a treatment, and then randomly sampling individuals without replacement. The shape of the IBR reflects only the shape of the underlying SAD 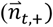.

It can be computationally intensive to compute rarefaction curves, and therefore analytical formulations of these curves are desirable to speed up computation. It is unlikely an analytical formulation of the sSBR exists because it requires averaging over each possible ordering of nearest sites; however, analytical expectations are available for the nsSBR and IBR. Specifically, we used the hypergeometic formulation provided by Hurlbert (1971) to estimate expected richness of the IBR. To estimate the nsSBR we extended Hurlbert’ s (1971) formulation (see Supplement S3). Our derivation demonstrates that the nsSBR is a rescaling of the IBR based upon the degree of difference in density between the two treatments under consideration. Specifically, we use the ratio of average community density to the density in the treatment of interest to rescale sampling effort in the individual based rarefaction curve. For a balanced design, the individual rarefaction curve of Treatment 1 can be adjusted for density effects by multiplying the sampling effort of interest by: (Σ_*t*_ Σ_*k*_,*N_t,k_*)/ (2 · Σ_*k*_ *N_1,k_*). Similarly, the Treatment 2 curve would be rescaled by (Σ_*t*_ Σ_*k*_ *N_t,k_*)/(2 · Σ_*k*_ *N_2,k_*). If the treatment of interest has the same density as the average community density then there is no density effect, and the nsSBR is equivalent to the IBR. Here we have based the density rescaling on average number of individuals, but alternatives exist, such as using maximum or minimum treatment density. Note that the nsSBR is only relevant in a treatment comparison that contrasts with the other two rarefaction curves, which can be constructed independently of any consideration of treatment effects.

### The mechanics of isolating the distinct effects of spatial aggregation, density, and SAD

The three curves capture different components of community structure that influence richness changes across scales (measured in number of samples or number of individuals, both of which can be easily converted to area, Table 1). Therefore, if we assume the components contribute additively to richness, then the effect of a treatment on richness propagated through a single component at any scale can be obtained by subtracting the rarefaction curves from each other. For simplicity and tractability, we assume additivity to capture first-order effects. This assumption is supported by Tjørve *et al*.’s (2008) demonstration that an additive partitioning of richness using rarefaction curves reveals random sampling and aggregation effects when using presence-absence data. We further validated this assumption using sensitivity analysis (see Supplement S5). Below we describe the algorithm to obtain the distinct effect of each component. Figure 5 provides a graphic illustration.

#### i) Effect of aggregation

The difference between the sSBRs of two treatments, 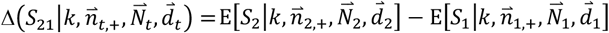, gives the observed difference in richness between treatments across scales (Fig. 5a, d, purple shaded area and solid curve respectively). It encapsulates the treatment effect propagated through all three components: shape of the SAD, density of individuals, and spatial aggregation. Differences between treatments in any of these factors could potentially translate into observed difference in species richness.

Similarly, the difference between the nsSBRs, 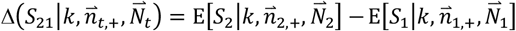, gives the expected difference in richness across treatments when spatial aggregation is removed (Fig. 5b, e, purple shaded area and dashed curve respectively). The distinct effect of aggregation across treatments from one plot to *k* plots can thus be obtained by taking the difference between the two Δ*S* values (Fig. 5g, i, green shaded area and solid line respectively), i.e.,

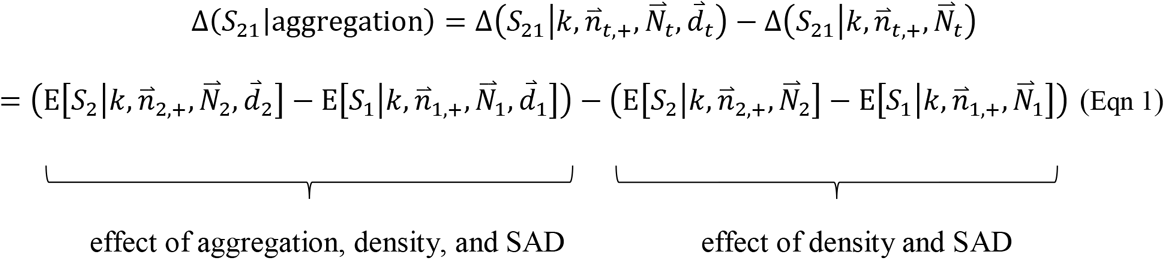

Equation 1 demonstrates that the effect of aggregation can be thought of as the difference between treatment effects quantified by the sample-based accumulation and sample-based rarefaction curves. An algebraic rearrangement of Eqn 1 demonstrates that Δ(*S*_21_|aggregation) can also be thought of as the difference between the treatments of the same type of rarefaction curve:

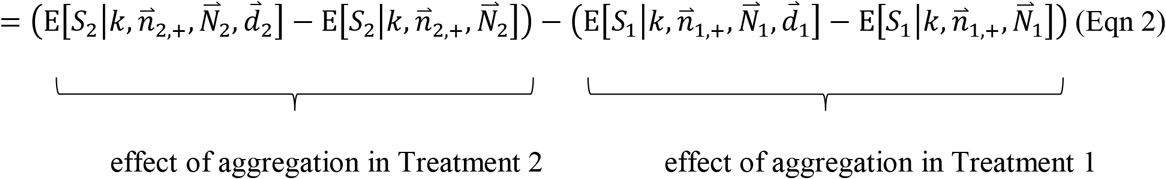

This simple duality can be extended to the estimation of the density and SAD effects, but we will only consider the approach laid out in Eqn 1 below. In Fig. 5, we separate each individual effect using the approach of Eqn 1 while the code in the `mobr` package uses the approach of Eqn 2.

One thing to note is that the effect of aggregation always converges to zero at the maximal spatial scale (*k* = *K* plots) for a balanced design. This is because, when all plots have been accumulated, 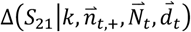 and 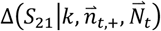 will both converge on the difference in total richness between the treatments. However, for an unbalanced design in which one treatment has more plots than the other, Δ(*S*_21_|aggregation) would converge to a nonzero constant because 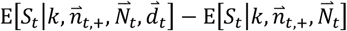 would be zero for one treatment but not the other at the maximal spatial scale (i.e., min(*K*_1_, *K*_2_) plots). This artefact is inevitable and should not be interpreted as a real decline in the relative importance of aggregation on richness, but simply as the diminishing ability to detect aggregation without sampling a larger region.

#### ii) Effect of density

In the same vein, the difference between the IBRs of the two treatments, 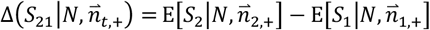, yields the treatment effect on richness propagated through the shape of the SAD alone, with the other two components removed (Fig. 5c, f, purple shaded area and dot-dashed curve respectively). The distinct effect of density across treatments from one individual to *N* individuals can thus be obtained by subtracting the Δ*S* value propagated through the shape of the SAD alone from the Δ*S* value propagated through the compound effect of the SAD and density (Fig. 5h, j, green shaded area and solid line respectively), i.e.,

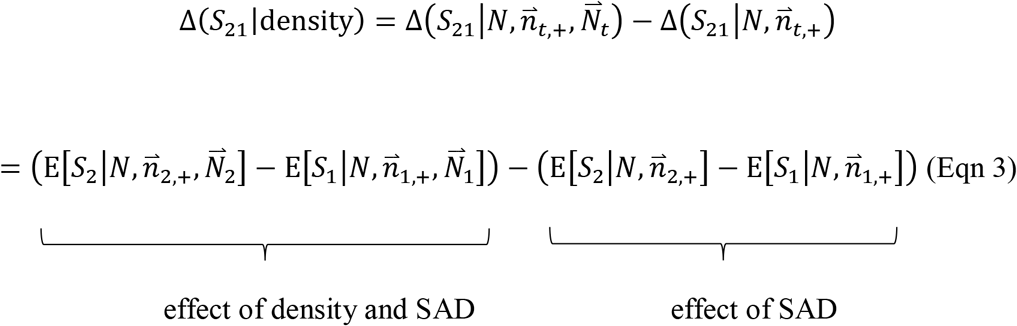

Note that in Eqn 3, spatial scale is defined with respect to numbers of individuals sampled (*N*) (and thus the grain size that would be needed to achieve this) rather than the number of samples (*k*).

#### iii) Effect of SAD

The distinct effect of the shape of the SAD on richness between the two treatments is simply the difference between the two IBRs (Fig. 5c, f, k, purple shaded area, dot-dashed curve, and green solid curve respectively), i.e.,

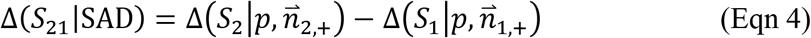

The scale of Δ(*S*_21_|SAD) ranges from one individual, where both individual rarefaction curves have one species and thus Δ(*S*_21_|SAD) = 0, to *N_min_* = min(*N*_1,+_, *N*_2, +_), which is the lower total abundance between the treatments.

The formulae used to identify the distinct effect of the three factors are summarized in Table 2.

**Table 2.** Calculation of effect size curves.

| Factor | Formula | Note |
| --- | --- | --- |
| Aggregation | $\Delta(S_{21}\vert \text{aggregation})$<br>$= \Delta(S_{21}\vert k, \vec{n}_{t,+}, \vec{N}_t, \vec{d}_t)$<br>$- \Delta(S_{21}\vert k, \vec{n}_{t,+}, \vec{N}_t)$ | Artificially, this effect always converges to zero at the maximal spatial scale ($K$ plots) for a balanced design, or a non-zero constant for an unbalanced design. |
| Density | $\Delta(S_{21}\vert \text{density})$<br>$= \Delta(S_{21}\vert N, \vec{n}_{t,+}, \vec{N}_t)$<br>$- \Delta(S_{21}\vert N, \vec{n}_{t,+})$ | To compute this quantity, the x-axes of the nsSBR are converted from plots to individuals using average individual density |
| SAD | $\Delta(S_{21}\vert SAD)$<br>$= \Delta(S_2\vert N, \vec{n}_{2,+})$<br>$- \Delta(S_1\vert N, \vec{n}_{1,+})$ | This is estimated directly by comparing the IBRs between two treatments. |

### Significance tests and acceptance intervals

In the multi-scale analysis, we also applied Monte Carlo permutation procedures to 1) construct acceptance intervals (or non-rejection intervals) across scales on simulated null changes in richness, and 2) carry out goodness of fit tests on each component (Loosmore and Ford 2006, Diggle-Cressie-Loosmore-Ford test [DCLF]; Baddeley et al. 2014). See Supplement S4 for descriptions of how each set of randomizations was developed to generate 95% acceptance intervals (Δ*S*_null_), which can be compared to the observed changes (Δ*S*_obs_). Strict interpretations of significance in relation to the acceptance intervals is not warranted because each point along the spatial scale (x-axis) is effectively a separate comparison. Consequently, a problem arises with multiple non-independent tests and the 95% bands cannot be used for formal significance testing due to Type I errors. The DCLF test (see Supplement S4) provides an overall significance test with a proper Type I error rate (Loosmore and Ford, 2006), but this test in turn suffers from Type II error (Baddeley et al. 2014). There is no mathematical resolution to this and user judgement should be emphasized if formal *p*-values are needed.

### Sensitivity Analysis

Although the logic justifying the examination separating the effect of the three components is rigorous, we tested the validity of our approach (and the significance tests) by simulations using the R package `mobsim` (May et al. 2018). The goal was to establish the rate of type I error (i.e., detecting significant treatment effect through a component when it does not differ between treatments) and type II error (i.e., nonsignificant treatment effect through a component when it does differ). This was achieved by systematically comparing simulated communities in which we altered one or more components while keeping the others unchanged (see Supplement S5). Overall, the benchmark performance of our method was good. When a factor did not differ between treatments, the detection of significant difference was low (Supplemental Table S5.1). Conversely, when a factor did differ, the detection of significant difference was high, but decreased at smaller effect sizes. Thus, we were able to control both Type I and Type II errors at reasonable levels. In addition, there did not seem to be strong interactions among the components – the error rates remained consistently low even when two or three components were changed simultaneously. The code to carry out the sensitivity analysis is archived here: https://github.com/MoBiodiv/mobr/blob/master/scripts/mobr_sensitivity.R

### An empirical example

In this section, we illustrate the potential of our method with an empirical example presented in Powell et al. (2013). Invasion of an exotic shrub, *Lonicera maackii*, has caused significant, but strongly scale-dependent, decline in the diversity of understory plants in eastern Missouri (Powell et al. 2013). Specifically, Powell et al. (2013) showed that the effect size of the invasive plant on herbaceous plant species richness was relatively large at plot-level spatial scales (1 m^2^), but the proportional effect declines with increasing windows of observations, with the effect becoming negligible at the largest spatial scale (500 m^2^). Using a null model approach, the authors further identified that the negative effect of invasion was mainly due to the decline in plant density observed in invaded plots. To recreate these analyses run the R code archived here: https://github.com/MoBiodiv/mobr/blob/master/scripts/methods_ms_figures.R.

The original study examined the effect of invasion across scales using the slope and intercept of the species-area relationship. We now apply our MoB approach to data from one of their sites from Missouri, where the numbers of individuals of each species were recorded from 50 1-m^2^ plots sampled from within a 500-m^2^ region in the invaded part of the forest, and another 50 plots from within a 500-m^2^ region in an adjacent uninvaded part of the forest. Our method leads to conclusions that are qualitatively similar to the original study, but with a richer analysis of the underlying components and their scale dependence. Moreover, our methods show that invasion influenced both the SAD and spatial aggregation, in addition to density, and that these effects went in opposite directions that depended on spatial scale.

Invasion decreased total abundance (*N*, Fig. 3, *p* = 0.001), suggesting a possible influence on *S*. Indeed, the two-scale analysis suggests that invasion decreases average richness (*S*) at the *α* (Fig. 4a, 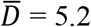, *p* = 0.001) but not *γ* scale (Fig. 4c, 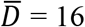, *p* = 0.438). Comparing rarefied richness (*S_n_*, Fig. 4d,f) allows us to test if the negative influence of invasion on *N* influenced *S* directly. Specifically, we found *S_n_* was higher in the invaded areas (significantly so 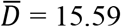, *p* = 0.001 at the *γ* scale evaluated at *n* = 250; Fig. 4f) which indicates that once the negative effect on abundance was controlled, invasion actually increased diversity through an increase in species evenness.

**Figure 3.**
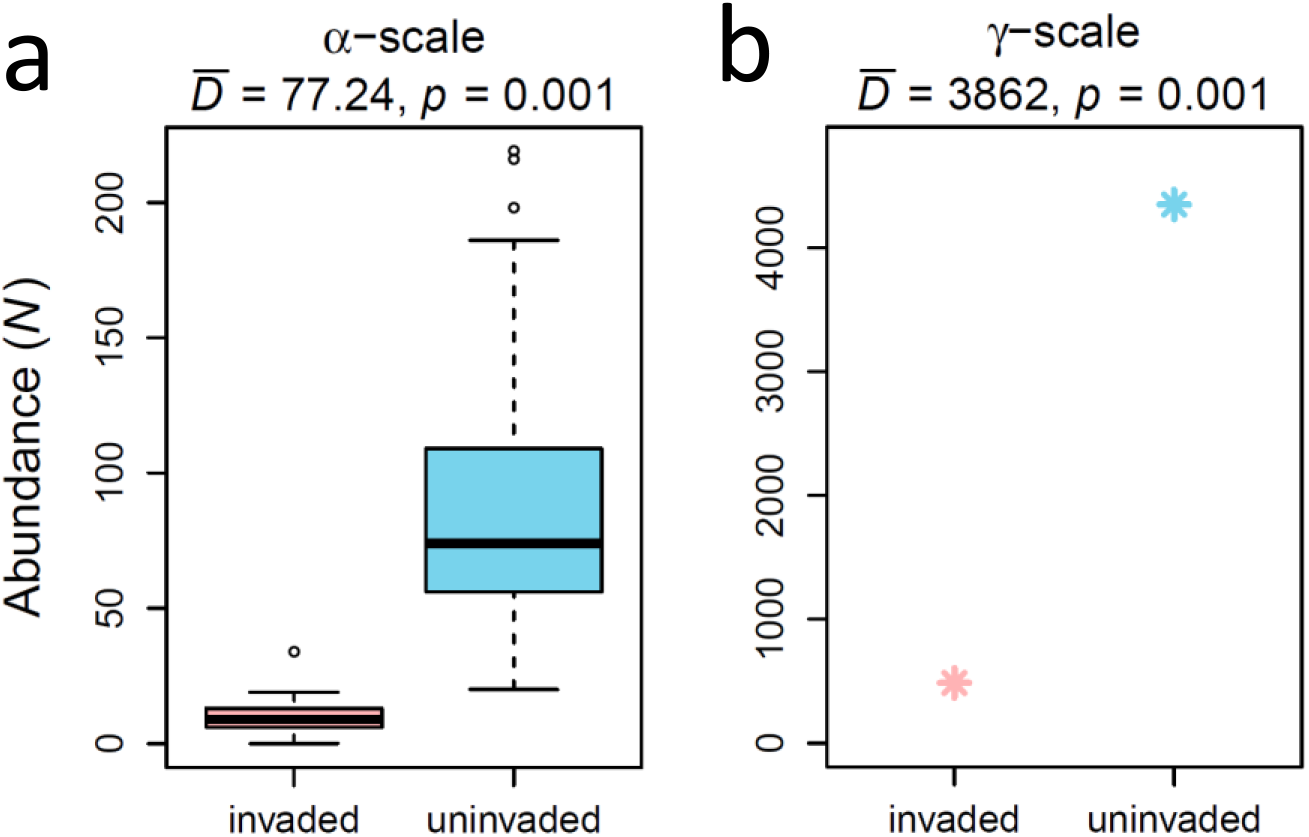
The total abundance (N) of vascular plant species in plots invaded (red boxplots and points) and uninvaded (blue boxplots and points) by Lonicera maackii at the *α*-scale (a, single plot) and the *γ*-scale (b, all plots). The p-values are based on 999 permutations of the treatment labels.

**Figure 4.**
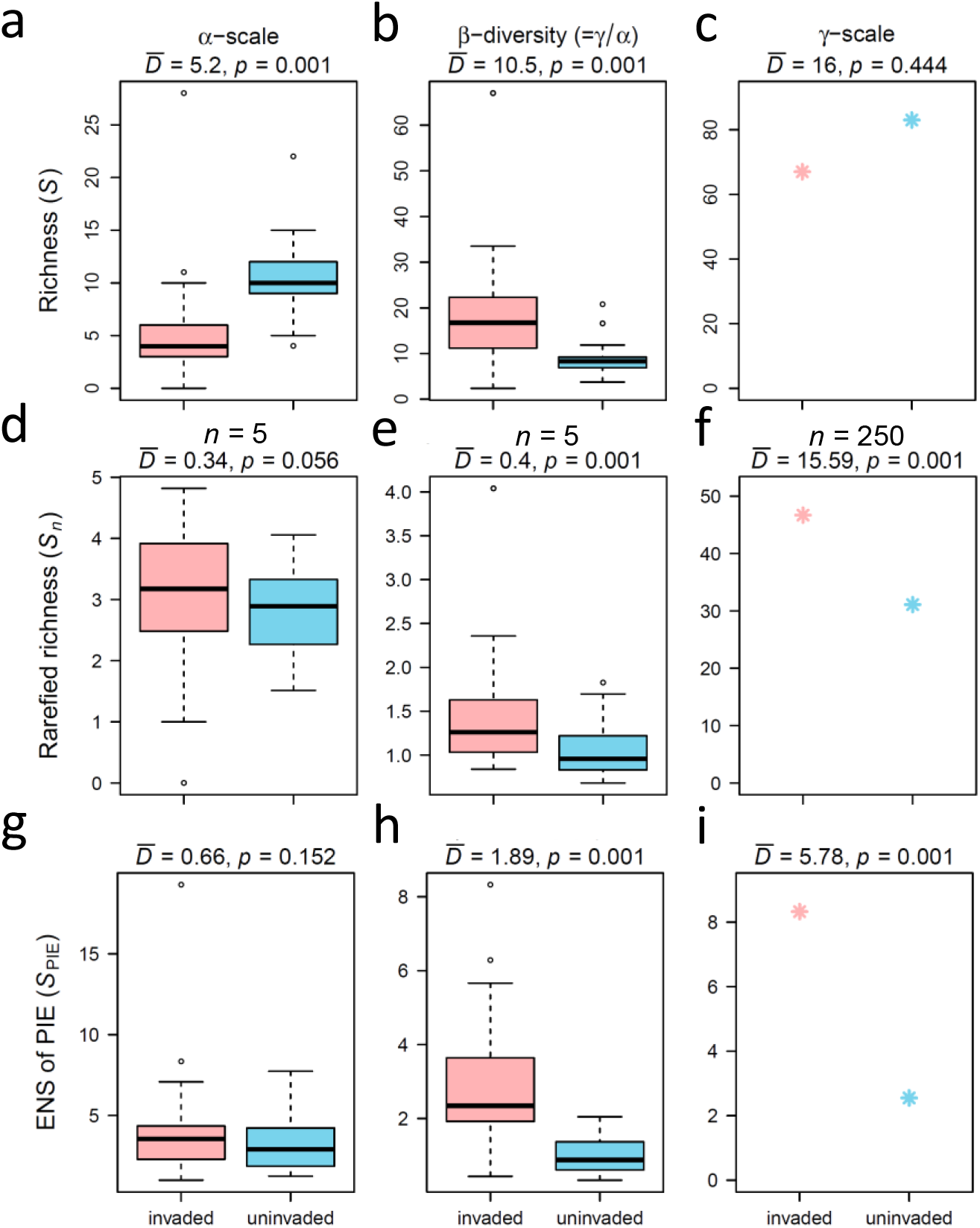
The two-scale analysis for the biodiversity metrics of the invaded (red boxplots and points) and uninvaded (blue boxplots and points) treatments at the *α* (i.e., single plot), *β* (i.e., between plots), and *γ* (i.e., all plots) scales. Rarefied richness (*S_n_*, panels d-f) were computed for *n* = 5, 5, and 250 individuals for the *α* (d), *β* (e), and *γ* (f) scales respectively. The *p*-values are based on 999 permutations of the treatment labels.

To identify whether the increase in evenness due to invasion was primarily because of shifts in common or rare species, we examined ENS of PIE (*S*_PIE_) which is more sensitive to common species relative to comparisons of *S*, which is more sensitive to rare species. At the *α* scale, invasion did not strongly influence *S*_PIE_ (Fig. 4g), but at the *γ* scale, there was evidence that invaded sites had greater evenness in the common species (Fig. 4i, 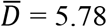, *p* = 0.001). In other words, the degree of dominance by any one species was reduced in the invaded sites.

The *β* diversity metrics were significantly higher (Fig. 4b,e,h *p* = 0.001) in the invaded sites, suggesting that invaded sites had greater spatial species turnover and thus were more heterogeneous. These increases in spatial turnover appeared to be only slightly due to the sole effect of increased spatial intraspecific aggregation in invaded sites as *β_Sn_* displayed the most modest effect size 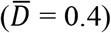. Therefore, it appears that the shift in SAD and decreased *N* also are playing a role in increasing *β*-diversity in the invaded treatment as they are reflected in *β_S_* and *β*_*S*_PIE__.

Overall the two-scale analysis indicates: 1) that there are scale-dependent shifts in richness, 2) that these are caused by invasion decreasing *N*, increasing evenness in common species, and increasing species patchiness.

Applying the multi-scale analysis, we further disentangled the effect of invasion on diversity through the three components (SAD, density, and aggregation) across all scales of interest. Figure 5a-c present the three sets of curves for the two treatments: the sSBR, in which plots accumulate by their spatial proximity (Fig. 5a); the nsSBR, in which individuals are randomized across plots within a treatment (Fig. 5b); and the IBR, in which species richness is plotted against number of individuals (Fig. 5c). Figure 5d-f show the effect of invasion on richness, obtained by subtracting the red curve from the blue curve for each pair of curves. The bottom panels (i-k), show the effect of invasion on richness through each of the three factors, is obtained by subtracting the curves in panels g and h from each other. The contribution of each component to difference in richness between the invaded and uninvaded sites is further illustrated in Fig. 5.

**Figure 5.**
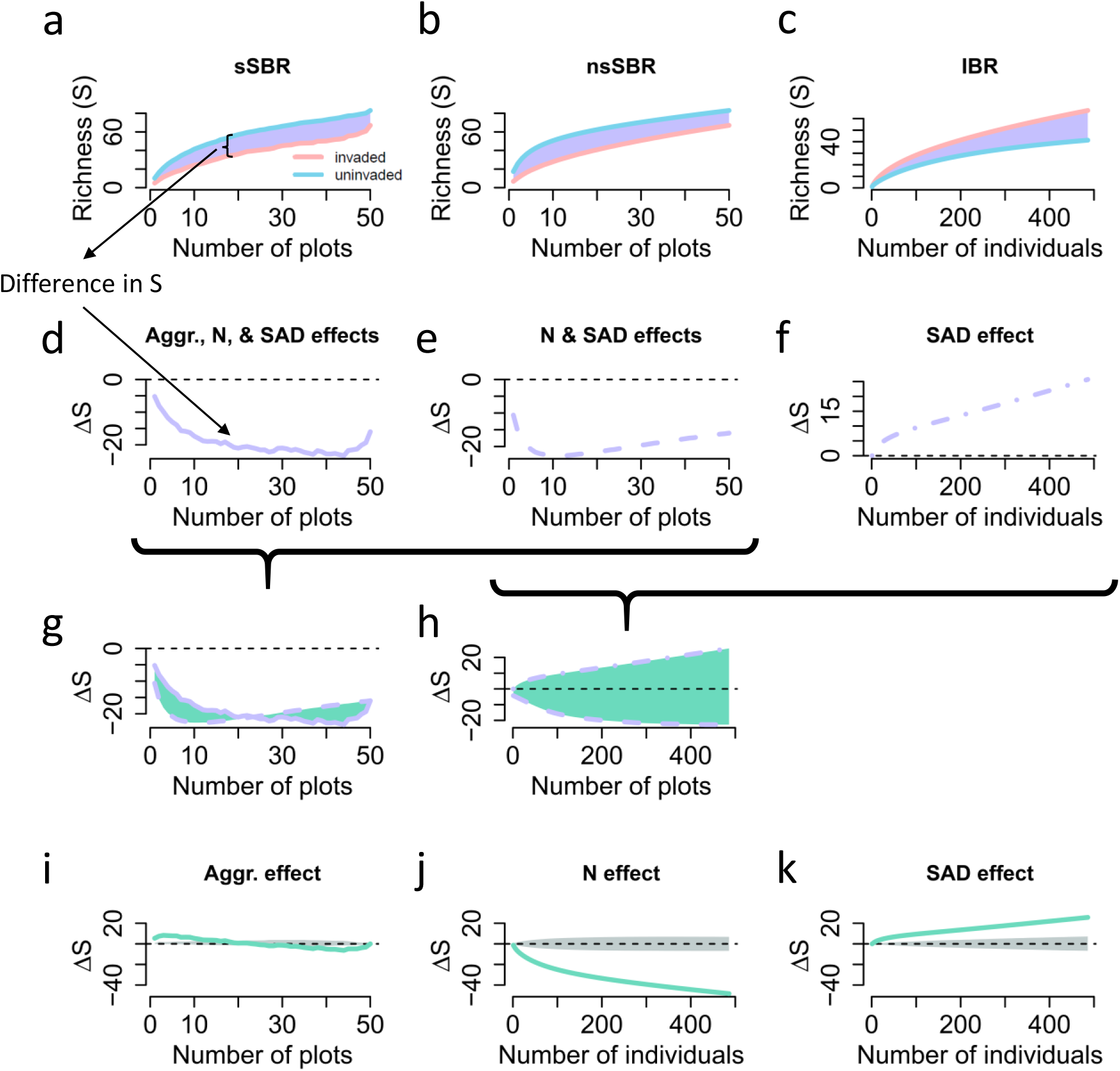
Application of the MoB multi-scale analysis to the invasion data set. Panels a-c, show the invaded (pink) and uninvaded (light-blue) in the three types of rarefaction curves (defined in Fig. 2 and Table 1). The purple polygons represent the difference (i.e., treatment effect) for each set of curves which is plotted in panels d-f. By taking the difference again (green shaded area and curves in g and h) we can obtain the treatment effect on richness through a single component (i-k). See text for details (Eqn 1.). The grey shaded area in i-k shows the 95% acceptance interval for each null model, the cross scale DCLF test for each factor was significant (*p* = 0.001). The dashed line shows the point of no-change in richness between the treatments. Note that in panel c) the IBR are only shown across their common range of individuals.

Consistent with the original study, our approach shows that the invaded site had lower richness than the uninvaded site at all scales (Fig. 5a). Separating the effect of invasion into the three components, we find that invasion actually had a positive effect on species richness through its impact on the shape of the SAD (Fig. 5k, Fig. 6a), which contributed to approximately 20% of the observed change in richness (Fig. 6b). This counterintuitive result suggests that invasion has made the local community more even, meaning that the dominant species were most significantly influenced by the invader. However, this positive effect was completely overshadowed by the negative effect of invasion on species richness through reductions in the density of individuals (Fig. 5j, Fig. 6a), which makes a much larger contribution to the effect of invasion on richness (as large as 80%, Fig. 6b). Thus, the most detrimental effect of invasion was the sharp decline in the number of individuals. The effect of aggregation (Fig. 5i), is much smaller compared with the other two components and was most important at small spatial scales. Our approach thus validates the findings in the original study but provides a more comprehensive way to quantify the contribution to richness decline caused by invasion by each of the three components, across spatial scales.

**Figure 6.**
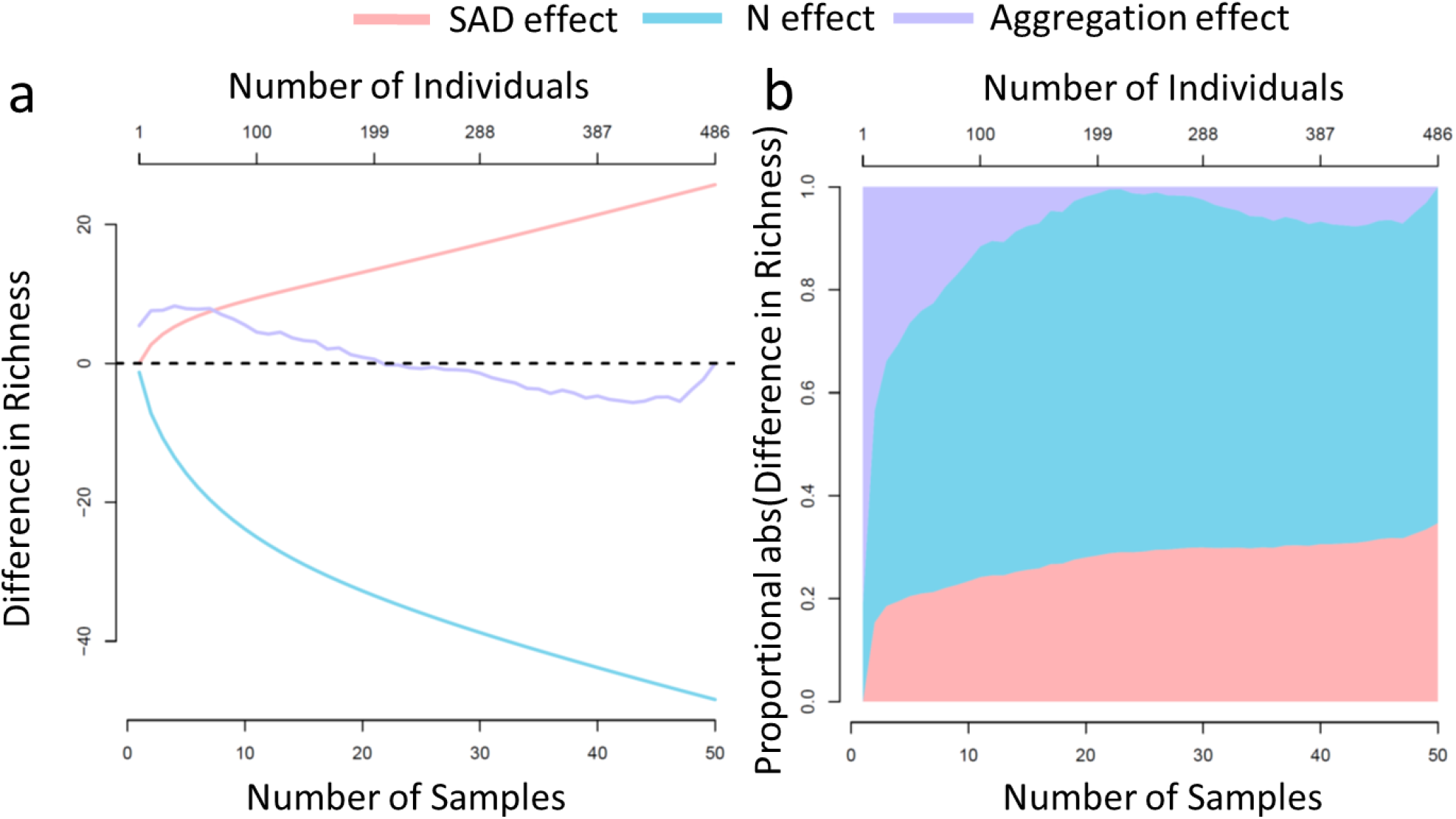
The effect of invasion on richness via individual effects on three components of community structure: SAD in red, density in blue, aggregation in purple across scales. The raw differences (a) and proportional stacked absolute values (b). The x-axis represents sampling effort in both numbers of samples (i.e., plots) and individuals (see top axis). The rescaling between numbers of individuals and plots we carried out by defining the maximum number of individuals rarefied to (486 individuals) as equivalent to the maximum number of plots rarefied to (50 plots), other methods of rescaling are possible. In panel (a) the dashed black line indicates no change in richness.

## Discussion

How does species richness differ between experimental conditions or among sites that differ in key parameters in an observational study? This fundamental question in ecology often lacks a simple answer, because the magnitude (and sometimes even the direction) of change in richness is strongly influenced by spatial scale (Chalcraft et al. 2004, Fridley et al. 2004, Knight and Reich 2005, Palmer et al. 2008, Powell et al. 2013, Blowes et al. 2017, Chase et al. 2018). Species richness is proximally determined by three underlying components—N, SAD and aggregation—which are also scale-dependent; this obscures the interpretation of the link between change in condition and change in species richness.

The MoB framework that we have introduced here provides a comprehensive answer to this question by taking a spatially explicit approach and decomposing the effect of the condition (treatment) on richness into its individual components. The two-scale analysis provides a big-picture understanding of the differences and components of richness by only examining the single plot (*α*) and all plots combined (*γ*) scales. The multi-scale analysis expands the endeavor to cover a continuum of scales, and quantitatively decomposes change in richness into three components: change in the shape of the SAD, change in individual density, and change in spatial aggregation. As such, we can not only quantify how richness changes at any scale of interest, but also identify how the change occurs and consequently push the ecological question to a more mechanistic level. For example, we can ask to what extent the effects on species richness are driven by numbers of individuals. Or instead, whether common and rare species, or their spatial distributions, are more strongly influenced by the treatments.

Here we considered the scenario of comparing a discrete treatment effect on species richness, but clearly the MoB framework will need to be extended to other kinds of experimental and observational designs and questions (Supplement S6). The highest priority extension of the framework is to generalize it from a comparison of discrete treatment variables to continuous drivers such as temperature and productivity. Additionally, we recognize that abundance is difficult to collect for many organisms and that there is a need to understand if alternative measures of commonness (e.g., visual cover, biomass) can also be used to gain similar insights.

Finally, we have only focused on taxonomic diversity here, whereas other types of biodiversity—most notably functional and phylogenetic diversity—are often of great interest, and comparisons such as those we have overviewed here would also be of great importance for these other biodiversity measures. Importantly, phylogenetic and functional diversity measures share many properties of taxonomic diversity that we have overviewed here (e.g., scale-dependence, non-linear accumulations, rarefactions, etc) (e.g., Chao et al. 2014), and it would seem quite useful to extend our framework to these sorts of diversities. We look forward to working with the community to develop extensions of the MoB framework that are most needed for understanding scale dependence in diversity change and overcome the limitations of the framework as currently implemented (Supplement S6).

MoB is a novel and robust approach that explicitly addresses the issue of scale-dependence in studies of diversity, and quantitatively disentangles diversity change into its three components. Our method demonstrates how spatially explicit community data and carefully framed comparisons can be combined to yield new insight into the underlying components of biodiversity. We hope the MoB framework will help ecologists move beyond single-scale analyses of simple and relatively uninformative metrics such as species richness alone. We view this as a critical step in reconciling much confusion and debate over the direction and magnitude of diversity responses to natural and anthropogenic drivers. Ultimately accurate predictions of biodiversity change will require knowledge of the relevant drivers and the spatial scales over which they are most relevant, which MoB (and its future extensions), helps to uncover.

## Acknowledgments

This paper emerged from several workshops funded with the support (to JMC) from the German Centre for Integrative Biodiversity Research (iDiv) Halle-Jena-Leipzig funded by the German Research Foundation (FZT 118) and by the Alexander von Humboldt Foundation as part of the Alexander von Humboldt Professorship of TMK. DJM was also supported by College of Charleston startup funding. We further thank N. Sanders, J. Belmaker, D. Storch, N. Baker, the anonymous reviewers and associate editor for discussions and comments on our approach.

## Data Accessibility

The data is archived with the R package on GitHub: https://github.com/mobiodiv/mobr

